# Benchmarking Hayai-Annotation Plants: A Re-evaluation Using Standard Evaluation Metrics

**DOI:** 10.1101/2023.09.08.556781

**Authors:** Andrea Ghelfi, Kenta Shirasawa, Sachiko Isobe

## Abstract

The rapid growth of next-generation sequencing (NGS) technology has led to a surge in the determination of whole genome sequences in plants. This has created a need for functional annotation of newly predicted gene sequences in the assembled genomes. To address this, “Hayai-Annotation Plants” was developed as a gene functional annotation tool for plant species. In this report, we compared Hayai-Annotation Plants with Blast2GO and TRAPID, focusing on the three primary gene-ontology (GO) domains: Biological Process (BP), Molecular Function (MF), and Cellular Component (CC). Using the *Arabidopsis thaliana* GO annotation as a benchmark, we evaluated each tool using two approaches: the area under the precision-recall curve (AUC-PR) and the metrics used at the critical assessment of functional annotation (CAFA). In the latter case, a CAFA-evaluator, was used to determine the F-score, weighted F-score, and S-score for each domain. Hayai-Annotation Plants showed better performances in all three GO domains. Our results thus reaffirm the effectiveness of Hayai-Annotation Plants for functional gene annotation in plant species. In this era of extensive whole genome sequencing, Hayai-Annotation Plants will serve as a valuable tool that facilitates simplified and accurate gene function annotation for numerous users, thereby making a significant contribution to plant research.

## 1. Introduction

The advancement of next-generation sequencing (NGS) technology has led to the construction of numerous whole genome sequences in plants as well as humans and other organisms. As a result, there is an increasing need for functional annotation for newly predicted gene sequences on the assembled genomes. Functional annotation of genes involves comparing query sequences with databases containing gene sequences and their functional information, such as OrthoDB (https://www.orthodb.org/), UniProt (https://www.uniprot.org/), and RefSeq (https://www.ncbi.nlm.nih.gov/refseq/). While referencing multiple databases enables more accurate functional annotation, the computational demands are high. This has made it challenging for many researchers who lack access to large-scale computer servers to perform high-precision functional annotation. In order to achieve fast and accurate gene function annotation, we have developed “Hayai-Annotation Plants,” an R package designed for the ultrafast and comprehensive functional annotation of plants (Ghelfi et al. 2018). Hayai-Annotation Plants employs the browser interface of the R package “Shiny”, alongside the cost-free edition of USEARCH (32-bit) (Edgar 2010) for sequence alignment. Hayai-Annotation Plants leverages UniProtKB data (Embryophyta taxonomy) to assign protein names, gene ontology (GO) terms and codes, EC numbers, protein existence levels, and evidence types. For obtaining GO codes, the Gene Ontology Annotation (UniProt-GOA) database was chosen due to its focus on delivering superior GO annotation quality (Camon et al. 2004). Details of the design of Hayai-Annotation Plants are given in our previous paper (Ghelfi et al. 2018), where the package was introduced.

It is worth noting that the remarkable speed of Hayai Annotation is largely due to its implementation of USEARCH, a renowned software package known for its fast and accurate sequence alignment capabilities. In our previous report (Ghelfi et al. 2018), we also evaluated and compared the accuracy of GO assignments annotated by Hayai-Annotation Plants and two other software programs: Blast2GO (Conesa and Götz 2008) and TRAPID (Van Bel et al. 2013). These software tools use different GO graph structures in annotation; Hayai-Annotation Plants and Blast2GO assign child GO terms, whereas TRAPID often reports a parental term rather than the most specific term (Van Bel and Vandepoele 2020).

Van Bel and Vandepoele (2020) pointed out that the comparison performed in our original paper on Hayai-Annotation Plants did not consider the GO ontology (Ashburner et al. 2000), even though GO ontology would have been particularly relevant to that analysis since the software tools analyzed use different levels of GO annotation. In consideration of this important limitation, we therefore re-evaluated the GO annotations of Hayai Annotation Plants by calculating the precision, recall and F_max_ for each GO domain independently (Clark and Radivojac 2013).

In the present study, we re-examined the GO assignments by Hayai-Annotation Plants in comparison to Blast2GO and TRAPID, with particular attention to the full spectrum of the GO ontology. Specifically, we focused on ontology terms with direct relationships within the three primary domains: Biological Process (BP), Molecular Function (MF), and Cellular Component (CC). By benchmarking against the gold standard GO annotation of Arabidopsis thaliana, we evaluated each software tool with two approaches: the area under the precision-recall curve (AUC-PR) and the standard metrics used in the critical assessment of functional annotation (CAFA). AUC-PR is a metric used to assess the performance of binary classification models, particularly in cases where the data is imbalanced (Sofaer et al. 2019). CAFA is a community-driven initiative that aims to assess and enhance the accuracy of computational methods used for predicting protein function (Zhou et al. 2019). We used CAFA-evaluator (https://github.com/BioComputingUP/CAFA-evaluator), which is the official tool for the CAFA challenges. Our results showed that Hayai-Annotation Plants exhibited better performance in each domain in this study.

## 2. Methods

### Data source

The GO annotation of A. thaliana downloaded from the Gene Ontology Consortium website (July 2019 from http://current.geneontology.org/products/pages/downloads.html) was considered the gold standard annotation.

### First method: Evaluation using AUC-PR

The evaluation AUC-PR was based on in-house scripts to assign the direct ontology of GO, which was obtained by extracting all relations defined as “is_a” in the go-basic obo file (downloaded in July 2019 from http://current.geneontology.org/ontology/subsets/goslim_plant.obo).

The sampling method employed the function “randomNumbers” from the R-package “random” (https://cran.r-project.org/web/packages/random/index.html) with 1,000 of the total number of annotated genes being considered for each replication. Three replications were conducted, and the mean values of each tool for each GO domain were compared based on the Tukey’s honestly significant difference (HSD) test at a significance level of 0.05.

The evaluation of GO annotation among Hayai-Annotation Plants, Blast2GO, and TRAPID was performed by calculating the AUC-PR for each one of the GO main domains (Molecular Function, Biological Process, and Cellular Component). The 1,000 genes comprising the sample were randomly selected from the annotated genes of Arabidopsis thaliana (Araport 11 representative peptide sequences), and three replications were considered for the comparisons. The complete methodology is available on GitHub (https://github.com/aghelfi/Hayai-Annotation-Plants/blob/master/evaluation/Ghelfi2023_Supplementary_Text_S1.pdf)

### Second method: Standard CAFA evaluation metrics

In order to compare the GO IDs predictions we calculated the F-score, weighted F-score and S-score and the precision-recall and remaining uncertainty-misinformation curves using the CAFA-evaluator (https://github.com/BioComputingUP/CAFA-evaluator). The software was executed using the same parameters as applied for the CAFA5 challenge (Kaggle; https://www.kaggle.com/competitions/cafa-5-protein-function-prediction), except that the “-no_orphans” parameter was added, which excludes the orphan term, for example the GO roots.

The gold standards file used in the CAFA-evaluator was generated by using an in-house python script to extract GO annotation from the go_arabidopsis.gaf file. The information accretion file was generated by using the Bioconductor SemDist (https://github.com/iangonzalez/SemDist) package and org.At.tair.db (https://bioconductor.org/packages/release/data/annotation/html/org.At.tair.db.html). The command lines and scripts implemented in this evaluation can be accessed on GitHub (https://github.com/aghelfi/Hayai-Annotation-Plants/tree/master/evaluation).

## Results

### First method: Evaluation using AUC-PR

The complete direct ontology was considered for the purpose of the calculations in all tested software tools, including the gold-standard GO annotation of *A. thaliana*. The results of Hayai-Annotation Plants and Blast2GO showed no significant differences at a significance level of *α*□=□0.05 in the MF and BP domains. TRAPID showed significantly lower values in all three GO domains compared to both Hayai-Annotation Plants and Blast2GO (Table 1).

**Table 1.**
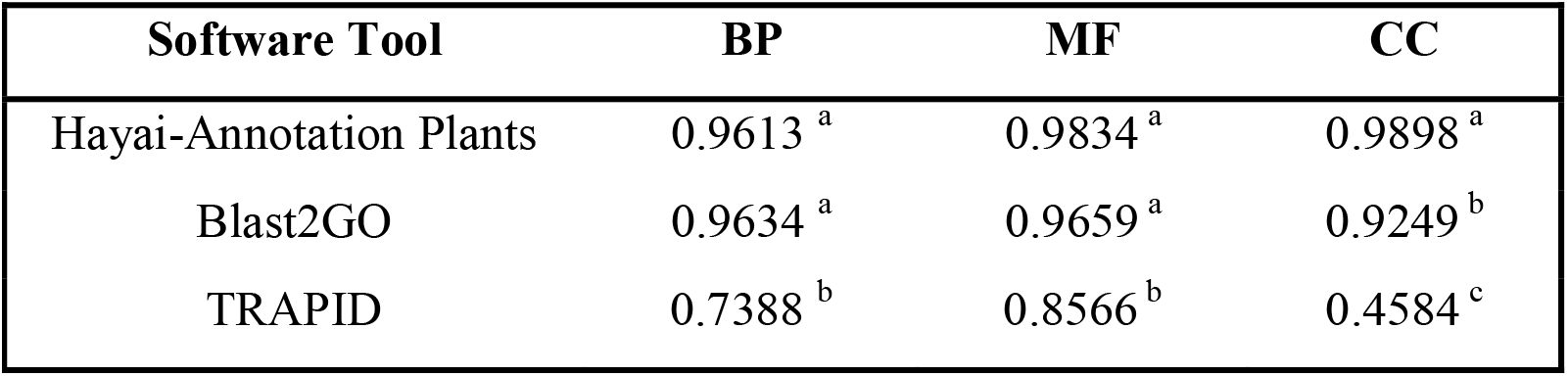
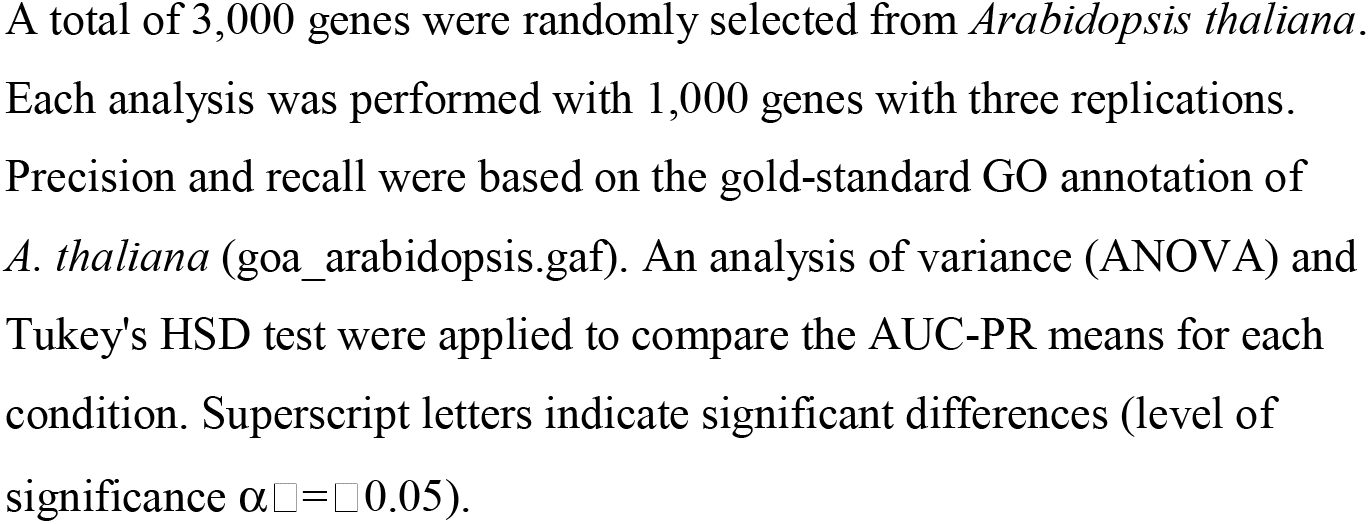
Average values from the overall performance evaluation using the AUC-PR to compare the different software tools.

| Software Tool | BP | MF | CC |
| --- | --- | --- | --- |
| Hayai-Annotation Plants | 0.9613<sup>a</sup> | 0.9834<sup>a</sup> | 0.9898<sup>a</sup> |
| Blast2GO | 0.9634<sup>a</sup> | 0.9659<sup>a</sup> | 0.9249<sup>b</sup> |
| TRAPID | 0.7388<sup>b</sup> | 0.8566<sup>b</sup> | 0.4584<sup>c</sup> |

A total of 3,000 genes were randomly selected from *Arabidopsis thaliana*. Each analysis was performed with 1,000 genes with three replications. Precision and recall were based on the gold-standard GO annotation of *A. thaliana* (goa\_arabidopsis.gaf). An analysis of variance (ANOVA) and Tukey's HSD test were applied to compare the AUC-PR means for each condition. Superscript letters indicate significant differences (level of significance $\alpha=0.05$ ).

Regarding the separation of test data versus training data, we divided 2,000 annotated genes of *Arabidopsis thaliana* (Araport11) into two groups of 1,000 genes each. We tested four values of sequence alignment in Hayai-Annotation Plants: 60%, 70%, 80%, and 90%. The results showed that, except for the CC domain, there was no significant difference among the different parameters selected for comparison (Table 2).

**Table 2.** Average values of the overall performance evaluation using AUC-PR for the comparison of different parameters of Hayai-Annotation Plants.

| Hayai-Annotation Plants | BP | MF | CC |
| --- | --- | --- | --- |
| SEQ_ID_90 | 0.9629<sup>a</sup> | 0.9875<sup>a</sup> | 0.9848<sup>a</sup> |
| SEQ_ID_80 | 0.9502<sup>a</sup> | 0.9822<sup>a</sup> | 0.9129<sup>c</sup> |
| SEQ_ID_70 | 0.9515<sup>a</sup> | 0.9837<sup>a</sup> | 0.9467<sup>b</sup> |
| SEQ_ID_60 | 0.9407<sup>a</sup> | 0.9711<sup>a</sup> | 0.9072<sup>c</sup> |

Four values of sequence alignment were compared: 60%, 70%, 80%, and 90%. An ANOVA and Tukey's HSD test were applied to compare the AUC-PR means for each condition. Superscript letters indicate significant differences (level of significance $\alpha=0.05$ ).

### Second method: Standard CAFA evaluation metrics

The same set of re-annotations of *A. thaliana* performed by Hayai-Annotation Plants and two benchmarks, BLAST2GO and TRAPID, was evaluated by using CAFA-evaluator. The GO annotation of *A. thaliana* from the Gene Ontology Consortium website was considered as a gold standard annotation for the purpose of our comparison. Using the CAFA-evaluator, three predictors were calculated for each GO domain independently, the F-score, weighted F-score and S-score, along with precision-recall and remaining uncertainty–misinformation curves as described in the CAFA. Hayai-Annotation Plants showed better performance when compared with our benchmarks in all GO domains in all calculated scores, as shown in Figure 1, Figure 2 and Figure 3.

**Figure 1.**
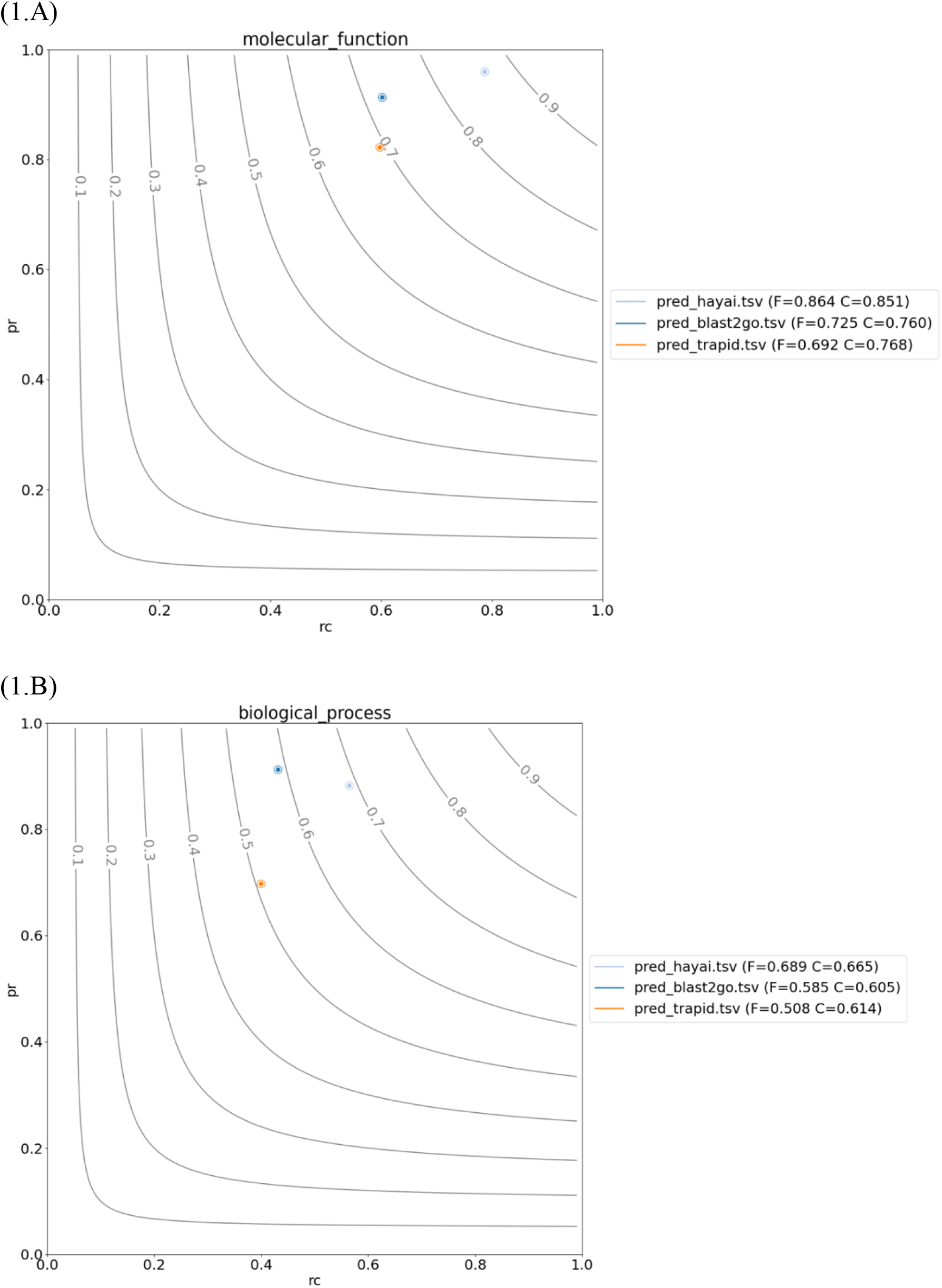
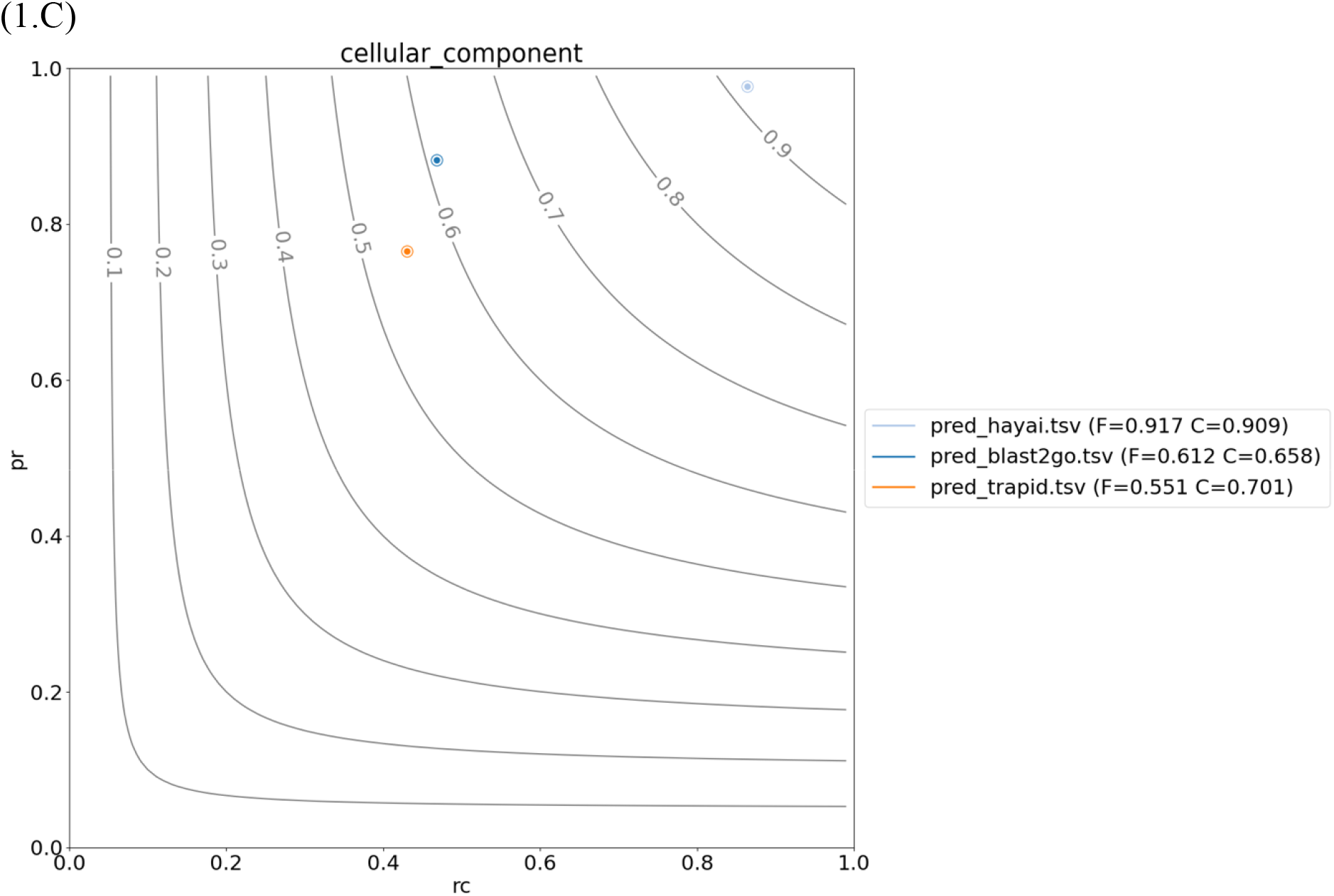
Fmax calculated based on precision (pr) and recall (rc) for each tool. (A) GO molecular function, (B) Biological process, (C) Cellular component.

**Figure 2.**
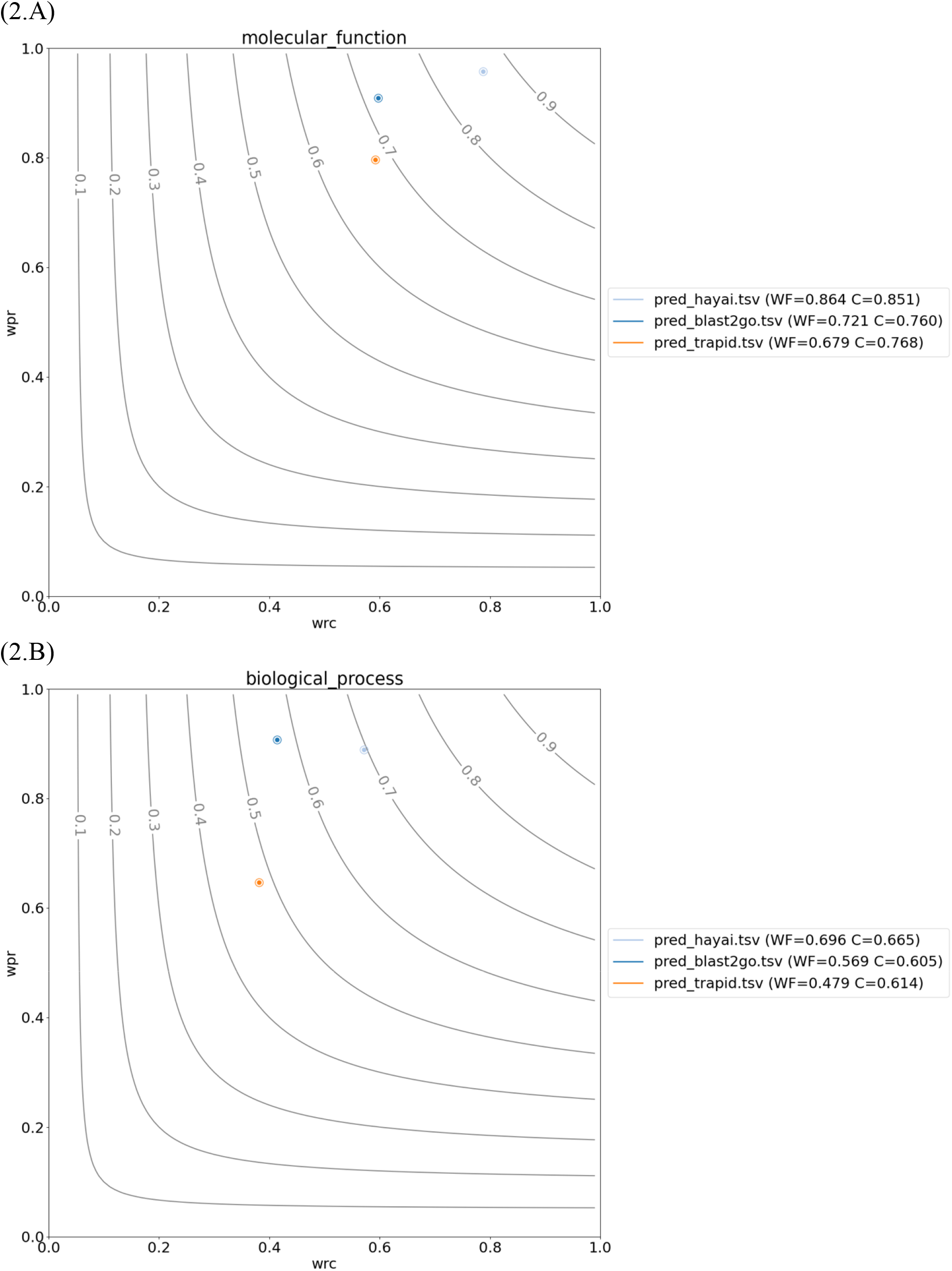
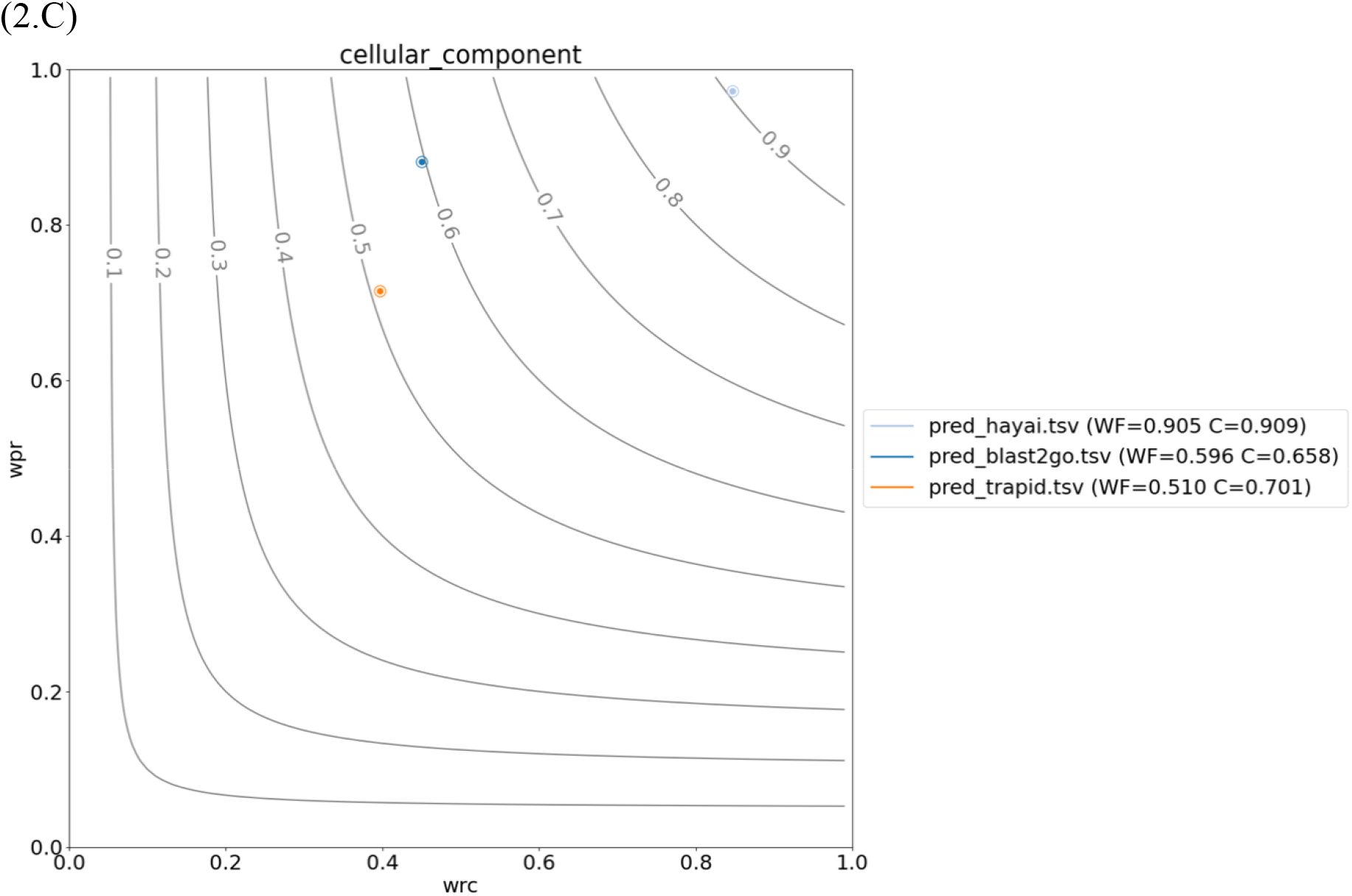
Fmax calculated based on weighted precision (wpr) and weighted recall (wrc) for each tool. (A) GO molecular function, (B) Biological process, (C) Cellular component.

**Figure 3.**
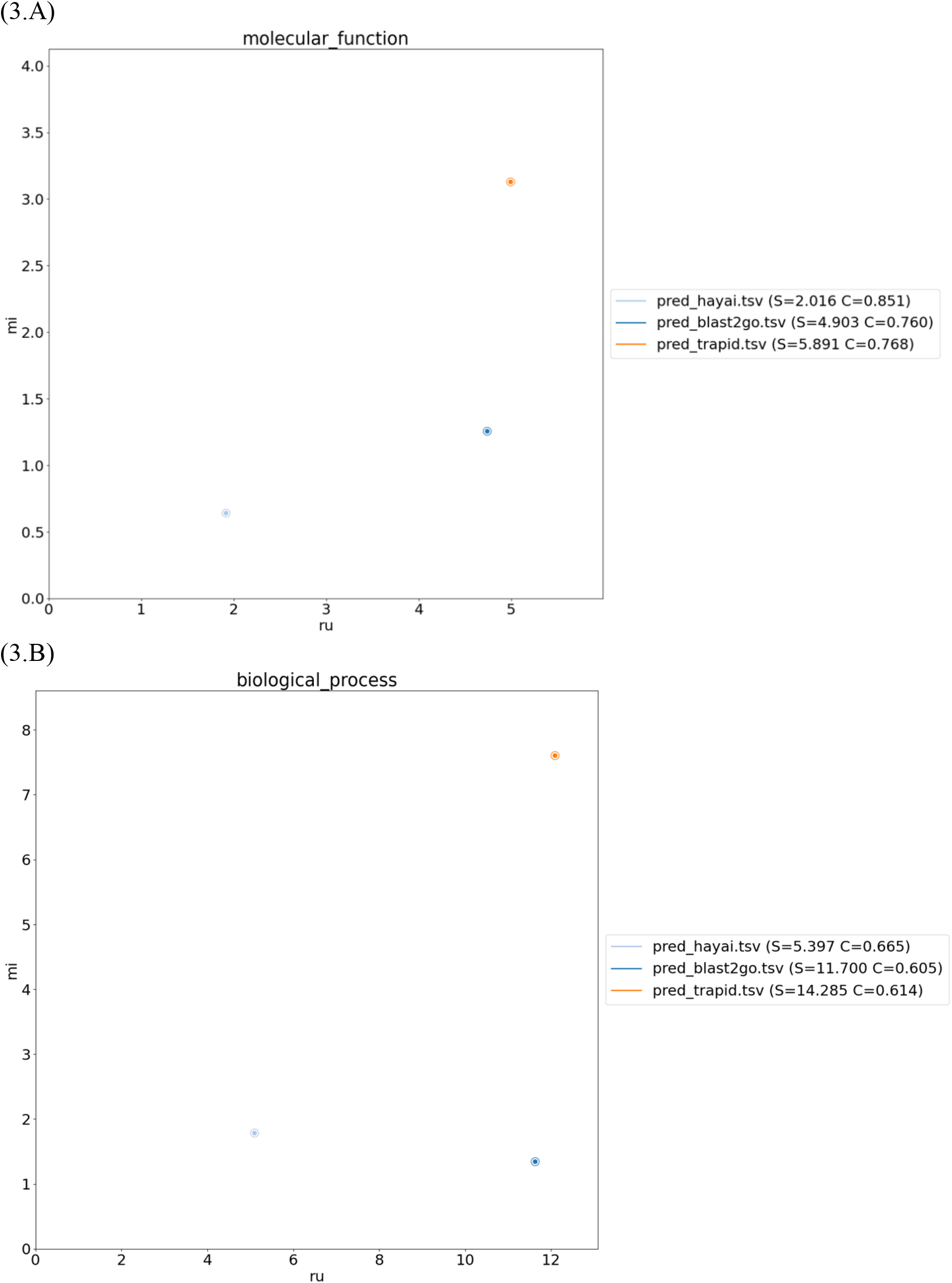
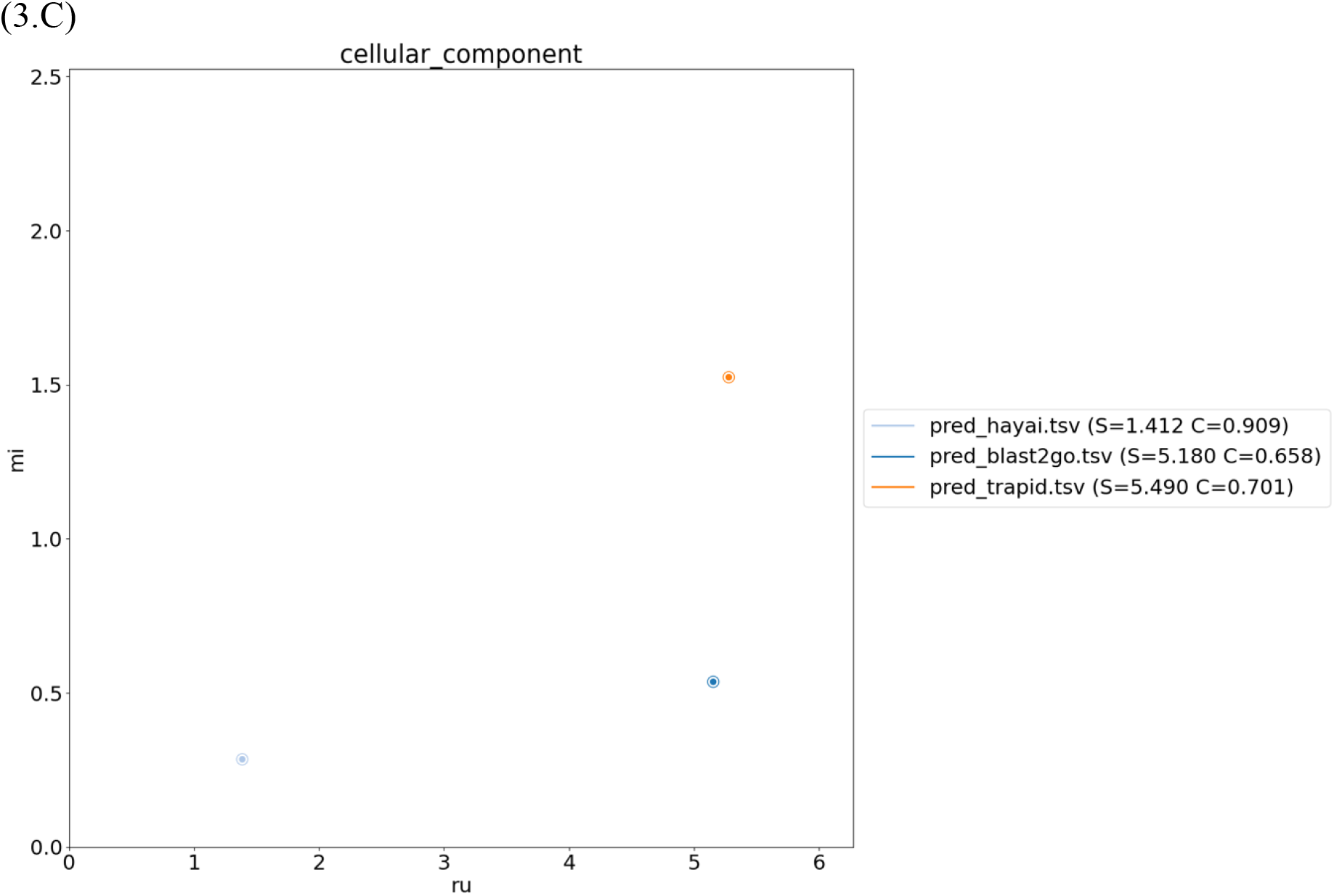
Semantic distance calculated by using remaining uncertainty (ru) and misinformation (mi) for each tool. (A) GO molecular function, (B) Biological process, (C) Cellular component.

## 4. Discussion and Conclusion

The results using AUC-PR showed that Hayai-Annotation Plants is a reliable tool to assign GO annotations, because it did not differ significantly (*α*□=□0.05) from the benchmark Blast2GO in all GO domains except for CC, in which Hayai-Annotation Plants showed better performances. The lower AUC-PR values observed with TRAPID were obtained because this tool annotates mainly the parental terms, and lower values would thus be expected compared to software tools that prioritize child GO terms, since we considered the complete GO set of information to calculate precision and recall.

In our previous publications about Hayai-Annotation Plants, we considered only child GO terms and disregarded parental terms to evaluate the performance of all tested software tools. In response to the observations by Van Bel and Vandepoele (2020), we realized that the comparison of tools in our previous publications was insufficient. Therefore, we utilized the AUC-PR comparison data described in this paper as a response to their comments and submitted it to the journal where the report of Van Bel and Vandepoele (2020) was published. Unfortunately, we were unable to secure the opportunity to publish the results of this analysis in the same journal where the comments were featured. As a result, despite our reservations about the situation, the paper commented upon by Van Bel and Vandepoele (2020) has been retracted.

Therefore, we considered it insufficient to rely solely on the evaluation using AUC-PR and conducted further assessment using the standard CAFA evaluation metric. The results again showed that Hayai-Annotation Plants is a reliable tool to assign GO annotations, because, in all GO domains, it showed better performance. In the review on gene functional annotation utilizing the Gene Ontology (GO) model, Zao et al. (2020) emphasize the significance of considering both the flat and hierarchical relationships among GO terms. In the context of Hayai-Annotation Plants, a more focused functional annotation is conducted by employing child terms from the GO to provide annotations. In contrast, TRAPID employs a higher level of GO terms for its annotations compared to Hayai-Annotation Plants. While comparing tools that utilize annotations from different levels of the GO hierarchy presents challenges, our re-evaluation, as mentioned earlier, has indeed validated the high reliability of functional annotation achieved by Hayai-Annotation Plants. Evaluating tools with different underlying systems in a definitive manner is exceedingly challenging, and the outcomes often vary based on the evaluation methods employed. What we aim to benchmark in this paper is not that the accuracy of TRAPID is inferior to that of Hayai-Annotation Plants, but rather that Hayai-Annotation Plants is a tool with high reliability and accuracy.

In essence, previous manuscripts concerning Hayai-Annotation Plants only carried out comparisons from a certain perspective, which might have raised issues if only the numerical data were extracted for comparison. However, this should not undermine the accuracy and reliability of Hayai-Annotation Plants as a functional annotation tool. The functionality of Hayai-Annotation Plants is publicly available through the following site (https://github.com/aghelfi/Hayai-Annotation-Plants), and analysis on the cloud is also feasible at http://pgdbjsnp.kazusa.or.jp/app/hayai2. Moreover, the Plant GARDEN platform has integrated the results of gene sequence re-annotation using a customized version of Hayai-Annotation Plants, further highlighting its capabilities (Ichihara et al. 2023). In the present era of extensive whole genome sequencing, Hayai-Annotation Plants will serve as a valuable tool that facilitates simplified and accurate gene function annotation for numerous users and will thereby make a significant contribution to plant research.

## References

Ashburner M, Ball CA, Blake JA, et al (2000) Gene Ontology: tool for the unification of biology. Nat Genet 25:25–29

Camon E, Magrane M, Barrell D, et al (2004) The Gene Ontology Annotation (GOA) Database: sharing knowledge in Uniprot with Gene Ontology. Nucleic Acids Res 32:D262–6

Clark WT, Radivojac P (2013) Information-theoretic evaluation of predicted ontological annotations. Bioinformatics 29:i53–61

Conesa A, Götz S (2008) Blast2GO: A comprehensive suite for functional analysis in plant genomics. Int J Plant Genomics 2008:619832

Edgar RC (2010) Search and clustering orders of magnitude faster than BLAST. Bioinformatics 26:2460–2461

Ghelfi A, Shirasawa K, Hirakawa H, Isobe S (2018) Hayai-Annotation Plants: an ultra-fast and comprehensive gene annotation system in plants. bioRxiv 473488

Ichihara H, Yamada M, Kohara M, et al (2023) Plant GARDEN: a portal website for cross-searching between different types of genomic and genetic resources in a wide variety of plant species. BMC Plant Biol 23:391

Sofaer HR, Hoeting JA, Jarnevich CS (2019) The area under the precision□recall curve as a performance metric for rare binary events. Methods Ecol Evol 10:565–577

Van Bel M, Proost S, Van Neste C, et al (2013) TRAPID: an efficient online tool for the functional and comparative analysis of de novoRNA-Seq transcriptomes. Genome Biol 14:1–10

Van Bel M, Vandepoele K (2020) Comment on ‘Hayai-Annotation Plants: an ultrafast and comprehensive functional gene annotation system in plants’: the importance of taking the GO graph structure into account. Bioinformatics 36:5558–5560

Zhou N, Jiang Y, Bergquist TR, et al (2019) The CAFA challenge reports improved protein function prediction and new functional annotations for hundreds of genes through experimental screens. Genome Biol 20:244

